# Developmental myelination is modified by microglial pruning

**DOI:** 10.1101/659482

**Authors:** Alexandria N. Hughes, Bruce Appel

## Abstract

During development, oligodendrocytes contact and wrap neuronal axons with myelin. Similar to neurons and synapses, excess myelin sheaths are produced and selectively eliminated. However, unlike these other structures, almost nothing is known about myelin sheath elimination. Microglia, the resident immune cells of the CNS, refine the developing CNS by engulfing surplus neurons and synapses. To determine if microglia also prune myelin sheaths, we used zebrafish to visualize and manipulate interactions between microglia, oligodendrocytes, and neurons during development. We found that microglia closely associate with oligodendrocytes and specifically phagocytose myelin sheaths. Silencing neuronal activity with botulinum toxin (BoNT/B) increased myelin engulfment by microglia. Furthermore, oligodendrocytes maintained excessive myelin sheaths following microglial ablation. Our work reveals a neuronal activity-regulated role for microglia in regulating myelination by oligodendrocytes.

## Main Text

Neuronal axon conduction velocity is supported by myelin, a specialized, proteolipid-rich membrane. During development, oligodendroglial cells, or oligodendrocytes, generate numerous nascent myelin sheaths along axons. Neuronal activity promotes the formation and maturation of myelin sheaths (*1*–*6*), processes that are mechanistically reminiscent to synaptogenesis (*7*). Like synapses, myelin sheaths are pruned after formation (*8*). However, we know almost nothing about the mechanisms that underlie myelin pruning. Synapses in the developing CNS can be pruned by microglia, the resident immune cells of the CNS. Microglia locate and engulf weak synapses in an activity-regulated manner, leaving stronger synapses intact (*9*). Could microglia also prune myelin sheaths to refine myelination?

To visualize microglia, we established a transgenic line of zebrafish, *Tg(mpeg1.1:mVenus-CAAX)*, in which *mpeg1.1* regulatory DNA drives expression of the membrane-tethered mVenus yellow fluorescent protein in CNS microglia (*10*). Crossing this line to animals carrying *Tg(mbpa:mCherry-CAAX)*, a transgene that labels oligodendrocyte membrane with mCherry, permitted us to see microglia-oligodendrocyte interactions (Fig. 1A). In the spinal cord at 4 days post-fertilization (dpf), the time at which *Tg(mbpa:mCherry-CAAX)* expression becomes visible, we found that microglia resided in both the dorsal and ventral myelinated tracts. At this stage, microglia have few, mostly primary, processes (*11*). By identifying and counting the structures that individual microglial processes appeared to contact, we determined that most processes associated with myelin sheaths rather than oligodendrocyte cell bodies or unlabeled targets (Fig. 1B, C). To test whether these processes were static or continued to survey other structures, we performed timelapse imaging of microglia (Fig. 1D, D’). We found that microglia continually withdrew and extended new processes within myelinated tracts (Fig. 1E), consistent with the possibility that microglia actively survey a large number of myelin sheaths.

**Figure 1.**
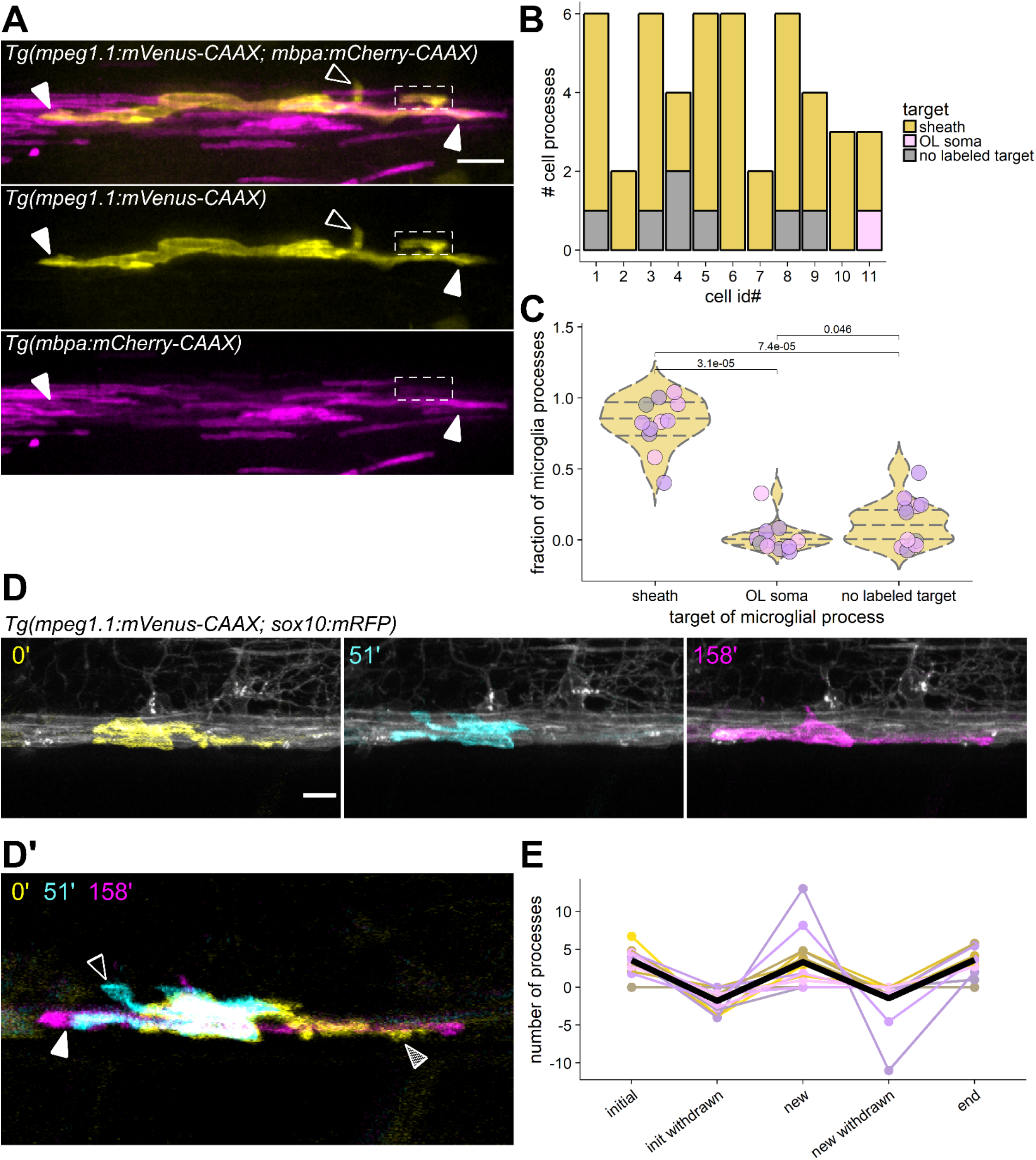
Microglia associate with and continuously survey myelin sheaths in the spinal cord. (A) A microglia (yellow) in the dorsal spinal cord of a *Tg(mpeg1.1:mVenus-CAAX; mbpa.mCherry-CAAX)* larva. Closed arrowheads mark close associations of microglial processes with myelin sheaths (magenta rectangles); open arrowheads mark processes extending toward putative unlabeled targets; dashed box shows a process enveloping a sheath. (B) Distribution of microglial processes per cell associated with myelin sheaths, oligodendrocyte somas, or unlabeled targets for n=11 cells (11 fish). (C) Fraction of microglial processes in (B) plotted per target type, Wilcox rank-sum test with Bonferroni-Holm correction for multiple comparisons. Each point represents one microglia; points are shaded so overlapping points are visible. (D, D’) Timelapse imaging frames of a microglia in a *Tg(mpeg1.1:mVenus-CAAX; sox10:mRFP)* larva, pseudocolored to show cell morphology at three different time points and merged in (D’). Striped arrowhead marks a process present at 0’ that is absent at 51’ (initial, withdrawn); open arrowhead marks a process generated during the imaging period that later disappears (new, withdrawn); closed arrowhead marks a process that is generated during the imaging period and is maintained at the end of the imaging window (new). (E) Number of processes generated and withdrawn by microglia during 1 hour of timelapse imaging; black trace is the mean (n=14 microglia in 14 fish). Scale bars, 10 μm.

Notably, our timelapse imaging experiments also captured occasional whole-sheath engulfment events by microglia (Fig. 2A, Movie S1). Following engulfment, myelin was only briefly visible within the microglia before fluorescence disappeared. We predicted that inhibiting the acidification of early phagosomes would delay the breakdown of phagocytosed myelin and allow us to determine how widespread myelin phagocytosis is among microglia (Fig. 2B). To inhibit phagosome acidification, we treated 4 dpf larvae with bafilomycin A1 for 1 hour prior to imaging, a procedure previously used in larval zebrafish to validate the pH-sensitivity of vital dyes (*12*). Microglia in *Tg(mpeg1.1:mVenus-CAAX; mbpa:mCherry-CAAX)* larvae treated with bafilomycin had more and brighter myelin inclusions than microglia of larvae treated with vehicle (Fig. 2C). We confirmed that myelin inclusions were contained within microglia by viewing cells in 3D (Movie S2). Morphological segmentation (see Methods) (*13*) allowed us to obtain measurements of these inclusions (Fig. 2D-H). We found that when larvae were treated with DMSO control, myelin inclusions were evident in about 2/3 of all microglia, with the remaining 1/3 containing no detectable inclusions. However, treatment with bafilomycin to acutely delay myelin breakdown revealed that all microglia contained myelin inclusions, suggesting that all spinal cord microglia participate in myelin pruning (Fig. 2E). Bafilomycin did not change the area of individual inclusions (Fig. 2F), but the total area of inclusions per microglia was increased (Fig. 2G), consistent with stalled myelin breakdown within microglia. Because oligodendrocytes can start myelinating at any time between 4 and 8 dpf, we also tested whether myelin phagocytosis detectable by bafilomycin might change over developmental time. We found no difference in the number of myelin inclusions per microglia at 4, 5, or 8 dpf, consistent with continuous myelin pruning by microglia over the course of developmental myelination (Fig. 2H).

**Figure 2.**
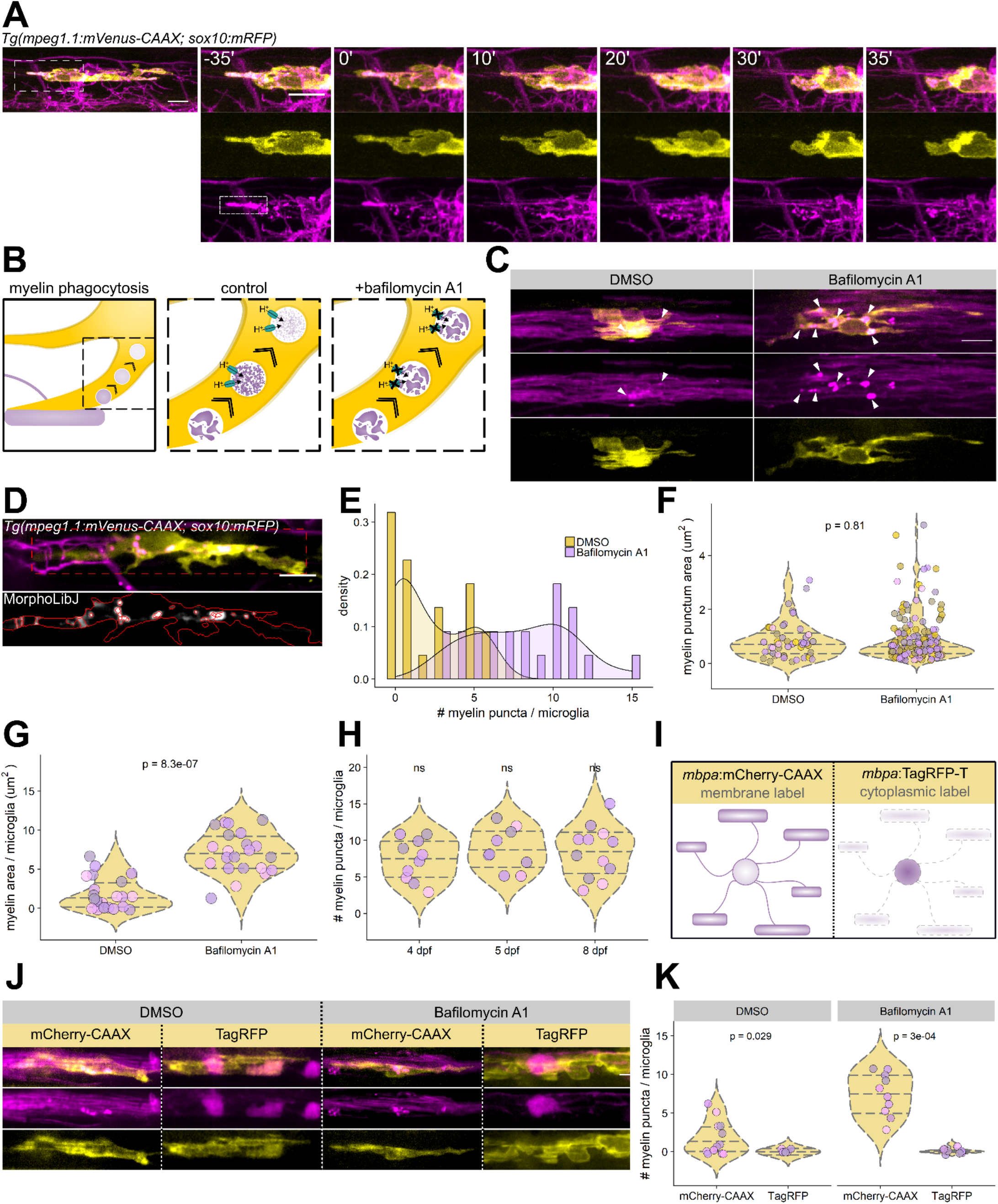
Microglia phagocytose myelin sheaths during developmental myelination. (A) Timelapse imaging of a microglia (yellow) interacting with an oligodendrocyte (magenta) in a *Tg(mpeg1.1:mVenus-CAAX; sox10:mRFP)* larva.Dashed box shows a myelin sheath (rectangle) that is engulfed and broken down by the microglia during the imaging period. (B) Strategy for delaying the breakdown of phagocytosed myelin with bafilomycin A1. (C) Microglia in *Tg(mpeg1.1:mVenus-CAAX; mbpa:mCherry-CAAX)* larvae treated with either DMSO or bafilomycin (1 μm) for 1 hour prior to imaging. Arrowheads mark round myelin inclusions inside microglia. (D) Example sum-projected image of a microglia containing mRFP inclusions (top) with morphological segmentation performed using the Fiji plugin MorphoLibJ to identify inclusions (bottom). (E, F, G) MorphoLibJ morphological segmentation on sum-projected images of microglia in 4 dpf *Tg(mpeg1.1:mVenus-CAAX; mbpa:mCherry-CAAX)* larvae that were treated with either DMSO or bafilomycin prior to imaging. Each point represents one inclusion (F) or microglia (G, H); points are shaded so overlapping points are visible (DMSO, n=22 cells from 22 larvae containing 48 inclusions; bafilomycin, n=22 cells/22 larvae/175 inclusions). (E) Histogram depicting the number of myelin inclusions per microglia in DMSO and bafilomycin-treated larva. (F) Area (μm^2^) of individual myelin inclusions inside microglia, Wilcox rank-sum test. (G) Total myelin inclusion area (μm^2^) per microglia, Wilcox rank-sum test. (H) Number of myelin inclusions per microglia in bafilomycin-treated larvae over the course of developmental myelination (4-8 dpf) (4 dpf, n=10 cells/10 fish/74 inclusions; 5 dpf, n=9 cells/9 fish/79 inclusions; 8 dpf, n=12 cells/12 fish/101 inclusions). (I) Schematic of transgenic oligodendrocyte reporters that label either oligodendrocyte membrane (mCherry-CAAX) or cytosol (TagRFP). (J) Examples of microglia in either *Tg(mpeg1.1:mVenus-CAAX; mbpa:mCherry-CAAX)* or *Tg(mpeg1.1:mVenus-CAAX; mbpa:TagRFP-T)* 4 dpf larvae, treated with either DMSO or bafilomycin prior to imaging. (K) Number of mCherry-CAAX and TagRFP inclusions per microglia in each of the four groups presented in (J), Wilcox rank-sum test with Bonferroni-Holm correction for multiple comparisons (mCherry-CAAX DMSO, n=12 cells/12 fish/18 inclusions; mCherry-CAAX bafilomycin, n=10 cells/10 fish/74 inclusions; TagRFP DMSO, n=6 cells/6 fish/0 inclusions; TagRFP bafilomycin, n=8 cells/8 fish/1 inclusion). Scale bars, 10 μm.

Although we only observed microglial phagocytosis of individual myelin sheaths, in principle, microglia might also engulf entire oligodendrocytes. To investigate this possibility, we repeated these experiments using a second oligodendrocyte reporter line, *Tg(mbpa:TagRFP-T)*, in which the same *mbpa* regulatory DNA drives expression of cytosolic TagRFP-T, a fluorophore with similar characteristics to mCherry *in vivo* (*14*). Because most oligodendrocyte cytosol is extruded from myelin sheaths, this line primarily labels oligodendrocyte cell bodies (Fig. 2I). We predicted that if microglia phagocytose entire oligodendrocytes then this oligodendrocyte somatic label would also be detectable within microglia. However, we very rarely detected TagRFP-T within microglia in DMSO or bafilomycin-treated larvae (Fig. 2J, K). These data suggest that microglia specifically engulf myelin sheaths without phagocytosing oligodendrocyte cell bodies. This specificity is consistent with previous work documenting an absence of oligodendrocyte lineage cell death during normal development in zebrafish (*15*).

Microglia contact numerous myelin sheaths (Fig. 1B, C) but do not phagocytose all sheaths. What directs microglia to phagocytose sheaths? Inhibiting neuronal activity increases synapse engulfment by microglia (*9*). To test whether microglial myelin phagocytosis is similarly regulated by neuronal activity, we mosaically silenced neurons by co-expressing botulinum toxin (BoNT/B) bicistronically with mTagBFP-CAAX, or labeled neurons with mTagBFP-CAAX alone, and used morphological segmentation to measure myelin phagocytosis by microglia at 4 dpf (Fig. 3A). In both the optic tectum and spinal cord, and with neurons both labeled and silenced, we found microglia that contained phagocytosed myelin (Fig. 3B). In the spinal cord, microglia can be found anywhere along the anterior-posterior axis, so we limited variability of myelin accessible to microglia by performing this analysis in optic tectum, a spatially-restricted region that contains more microglia than the spinal cord (11). We found that both myelin inclusion size and microglia size were unchanged by silencing neurons with BoNT/B (Fig. 3C, D). However, when neurons were silenced with BoNT/B, microglia contained many more myelin inclusions (Fig. 3E), and the total area of myelin inclusions was increased per microglia (Fig. 3F). Together, these data indicate that microglia prune more myelin when neuronal activity is silenced.

**Figure 3.**
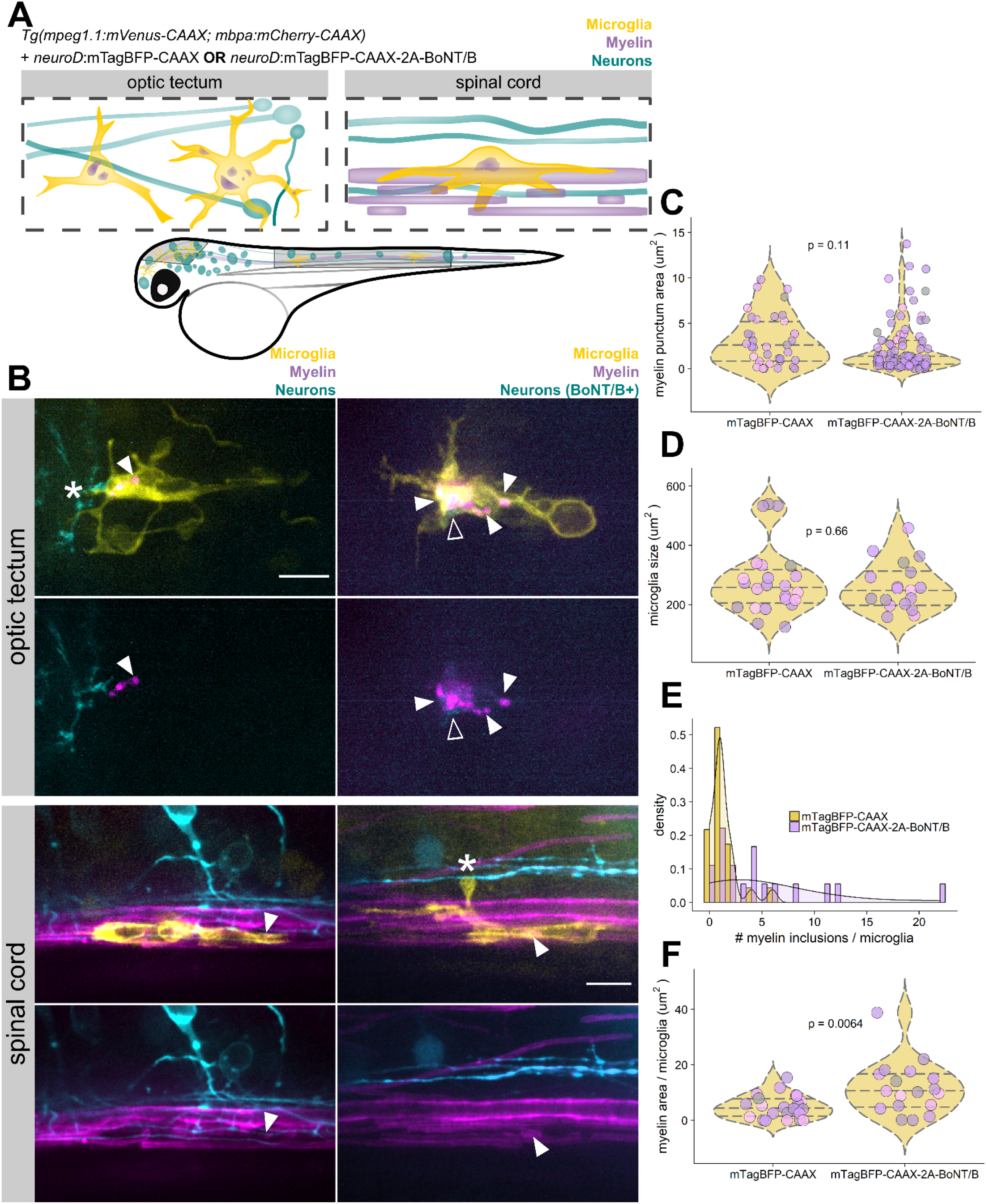
Microglial myelin phagocytosis is regulated by neuronal activity. (A) Schematic of transgenes used to label microglia and myelin (oligodendrocytes), and to label or manipulate neurons with botulinum toxin (BoNT/B), in both the optic tectum and spinal cord. (B) Microglia in the optic tectum (top panels) and spinal cord (bottom panels), in larvae carrying *Tg(mpeg1.1:mVenus-CAAX; mbpa:mCherry-CAAX)* to label microglia and myelin and mosaically expressing either neuroD:mTagBFP-CAAX (left) or neuroD:mTagBFP-CAAX-2A-BoNT/B (right) to label or silence neurons, respectively. Closed arrowheads mark myelin inclusions inside microglia (magenta round spots); open arrowheads mark neuronal inclusions (cyan round spots); asterisks mark microglial processes contacting neurons. Scale bars, 10 μm. (C, D, E, F) MorphoLibJ morphological segmentation on sum-projected images of optic tectum microglia in 4 dpf *Tg(mpeg1.1:mVenus-CAAX; mbpa:mCherry-CAAX)* larvae transiently expressing either mTagBFP-CAAX or mTagBFP-CAAX-2A-BoNT/B in neurons. Each point represents one inclusion (C) or microglia (D, F); points are shaded so overlapping points are visible (mTagBFP-CAAX, n=23 cells/18 fish/30 inclusions; mTagBFP-CAAX-2A-BoNT/B, n=18 cells/18 fish/87 inclusions). (C) Area (μm^2^) of individual myelin inclusions inside microglia, Wilcox rank-sum test. (D) Microglia area (μm^2^), Wilcox rank-sum test. (E) Histogram depicting the number of myelin inclusions inside microglia. (F) Total myelin inclusion area (μm^2^) per microglia, Wilcox rank-sum test.

Microglia eliminate some, but not all myelin sheaths, and microglia phagocytose more myelin when neuronal activity is blocked. Does microglial myelin phagocytosis direct myelin sheath refinement? To answer this, we generated a transgenic line to chemogenetically ablate microglia during myelination with the nitroreductase-metronidazole system. In this line, *Tg(mpeg1.1:NTR-2A-eGFP-CAAX)*, an enhanced variant of *E.coli* nitroreductase *(NfsB^T41Q/N71S/F124T^)* (NTR) (*16*) is expressed in microglia along with membrane-tethered eGFP. When larvae carrying this chemogenetic transgene were treated with 10 mM metronidazole for 24 hours microglia were ablated, but when larvae were treated with the vehicle control DMSO microglia survived and were labeled with eGFP (Fig. 4A). We quantified the amount of chemogenetic ablation of microglia by adapting a neutral red assay previously used for quantifying microglia (*17*, *18*) and determined that this strategy eliminates ~80% of microglia (Fig. 4B, C).

**Figure 4.**
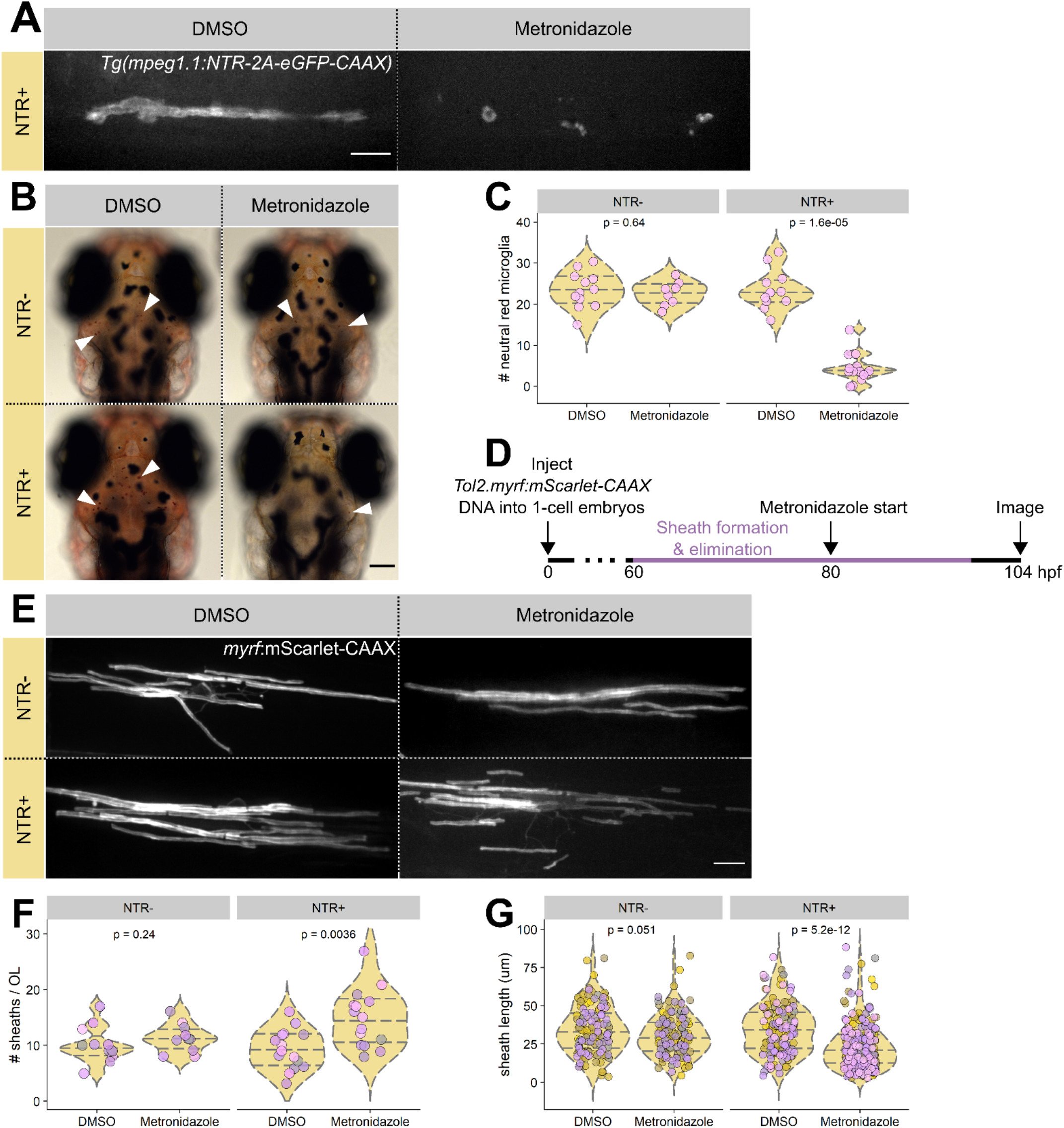
Microglial ablation during myelination increases myelin sheath number. (A) Microglia in *Tg(mpeg1.1:NTR-2A-eGFP-CAAX)* larvae treated with either DMSO vehicle or metronidazole (10 mM) for 24 hours prior to imaging. No microglia were present in the spinal cord of metronidazole treated, NTR+ larva, but fluorescent debris was visible as shown. Scale bar, 10 μm. (B) Brightfield images of neutral red-stained larvae in DMSO and metronidazole-treated NTR carriers (NTR+) and sibling noncarriers (NTR-). Arrowheads mark examples of neutral red-labeled microglia in optic tectum (small red dots). Scale bar, 100 μm. (C) Quantification of neutral red-labeled microglia in the optic tectum of larvaein each of the four groups. Each dot represents the neutral red cell count in one larva. Wilcox rank-sum with Bonferroni-Holm correction for multiple comparisons (NTR-DMSO, n=12 fish/281 microglia; NTR-metronidazole, n=8 fish/181 microglia; NTR+ DMSO, n=12 fish/283 microglia; NTR+ metronidazole, n=14 fish/63 microglia). (D) Strategy for labeling individual oligodendrocytes by microinjection at the 1-cell stage and chemogenetically ablating microglia with metronidazole in *Tg(mpeg1.1:NTR-2A-eGFP-CAAX)* larvae. (E) Example oligodendrocytes labeled by myrf:mScarlet-CAAX injected at 1-cell stage into *Tg(mpeg1.1:NTR-2A-eGFP-CAAX)* carriers (NTR+) and non-carrier siblings (NTR-) and treated with either DMSO or metronidazole at 80 hpf. Scale bar, 10 μm. (F, G) Quantification of sheath number (F) and length (G) for each of the four groups in (E) by tracing sheaths in 3D with Simple Neurite Tracer in Fiji. Each point represents one oligodendrocyte; points are shaded so overlapping points are visible. Wilcox rank-sum with Bonferroni-Holm correction for multiple comparisons (NTR-DMSO, n=12 cells/12 fish/121 sheaths; NTR-metronidazole, n=11 cells/11 fish/124 sheaths; NTR+ DMSO, n=15 cells/15 fish/139 sheaths; NTR+ metronidazole, n=15 cells/15 fish/223 sheaths). Scale bars, 10 μm.

We then used this microglia ablation approach to look at the myelin sheaths of individual oligodendrocytes. To do so, we performed microinjections of *Tol2.myrf:mScarlet-CAAX* DNA to scatter-label oligodendrocytes via Tol2-mediated transgenesis into 1-cell stage NTR+ embryos or sibling noncarriers (NTR-) (Fig. 4D). At 80 hpf, a time during peak myelin sheath formation, we administered metronidazole to eliminate microglia or DMSO control to both NTR carriers and noncarriers. At 104 hpf, we imaged oligodendrocytes in each of the four conditions (Fig. 4E). By tracing myelin sheaths formed by individual oligodendrocytes (*19*), we found that in the absence of microglia (NTR+, metronidazole) oligodendrocytes maintained a greater number of sheaths than oligodendrocytes in any of the three control groups (Fig. 4F). Intriguingly, these sheaths were also shorter in length (Fig. 4G), raising the possibility that sheaths normally pruned by microglia may be shorter than non-pruned sheaths. These data indicate that microglial sheath pruning contributes to normal developmental myelination by oligodendrocytes.

Our work identifies microglia as mediators of myelin refinement by pruning sheaths during development. Microglia often contain myelin debris in electron micrographs of diseased brain tissue (*20*, *21*), but our work suggests that myelin engulfment is a more widespread phenomenon that has likely evaded detection due to technical barriers to examining cell-cell interactions *in vivo*. Our data might help explain previous observations. Early work aimed at understanding myelin phagocytosis mechanisms in multiple sclerosis found that cultured macrophages engulfed healthy purified myelin (*22*). In a model of chemotherapy-induced cognitive impairment, microglial activation contributed to myelin thinning, which could be rescued by microglial ablation (*23*). Furthermore, scRNA-seq has detected oligodendrocyte-specific, myelin-enriched transcripts, such as *Mbp*, in microglia (*24*). In retrospect, these findings hinted that microglia phagocytose myelin in both health and disease. By revealing that microglia tune CNS myelination, our study now raises the possibility that microglia-oligodendrocyte interactions contribute to the development of brain function.

## Acknowledgments

We thank Drs. Marnie Preston and Jacob Hines for providing pME-NTR and p5E-myrf plasmids, respectively, and Dr. Wendy Macklin and Samantha Bromley-Coolidge for comments on the manuscript.

## Author contributions

A.N.H. and B.A. conceived the project. A.N.H. performed all the experiments and collected and analyzed all the data. A.N.H. wrote and B.A. edited the manuscript.

## Data and materials availability

All data are available in the manuscript or the supplementary materials. All plasmids and transgenic zebrafish are available by request.

## List of Supplementary Materials

Materials and Methods

Movies S1, S2

References (25–30)

## Materials and Methods

### Zebrafish lines and husbandry

All animal work was approved by the Institutional Animal Care and Use Committee at the University of Colorado School of Medicine. Zebrafish embryos were raised at 28.5°C in embryo medium and staged according to hours or days post fertilization (hpf/dpf) and morphological criteria (*25*).

We used the previously established transgenic lines *Tg(sox10:mRFP)^vu234^, Tg(mbpa:tagRFP)^co25^*, and *Tg(mbpa:mCherry-CAAX)^co13^*.

We also generated and used the new lines *Tg(mpeg1.1:mVenus-CAAX)^co56^* and *Tg(mpeg1.1:NTR-2A-eGFP-CAAX)^co57^*. All other reporters were expressed by transient transgenesis to achieve sparse labeling.

### Plasmid construction and generation of transgenic zebrafish

Tol2 expression plasmids were generated by Multisite Gateway cloning and injected into 1-cell embryos with Tol2 mRNA to generate transient transgenic animals. The following entry clones (*26*) were used in LR recombination reactions to generate expression plasmids:

p5E-neuroD, p5E-myrf, p5E-mpeg1.1

pME-NTR (*NfsB^T41Q/N71S/F124T^*), pME-mTagBFP-CAAX no stop, pME-mTagBFP-CAAX w/stop, pME-mVenus-CAAX, pME-mScarlet-CAAX

p3E-polyA, p3E-2A-BoNT/B, p3E-2A-eGFP-CAAX

pDEST-Tol2-pA, pDEST-Tol2-CG2 (green heart marker), pDEST-Tol2-CR2 (red heart marker)

### Drug treatments

Bafilomycin A1 (Tocris, cat. 1334) was diluted to 1 mM in DMSO and stored at −20°. To administer bafilomycin, we diluted this 1000x stock to 1 μM in embryo medium and treated larvae for 1 hour at 28.5° before mounting larvae in agarose for live imaging. An equivalent volume of DMSO was diluted 1000-fold in embryo medium (0.1%) as vehicle control.

Metronidazole (Sigma, M1547) was diluted to 10 mM in 0.1% DMSO in embryo medium immediately prior to use. Metronidazole is light sensitive, so all four treatment groups were kept in the incubator in the dark starting at administration (80 hpf) (*27*).

Neutral red (Sigma, N7005) was diluted to 2.5 μg/ml in embryo medium and administered to 4 dpf larvae for 2.5 hours at 28.5° in the dark (*17*, *18*). Following staining, larvae were washed 2-3 times in fresh media and monitored until nonspecific tissue redness washed out (30 min-1 hour) before mounting larvae dorsal-down in low-melt agarose for widefield, brightfield imaging (10x) of optic tectum microglia.

### Statistics

All statistics were performed in R (version 3.4.1) with RStudio. Plots were generated using dplyr and ggplot2 packages (28) with cowplot package formatting (29), and all statistical tests were performed using ggpubr (30) functions. We used the Wilcox rank-sum test (also called Mann-Whitney), with no assumption of normality, for all unpaired comparisons. For multiple comparisons, we first assessed global significance using the Kruskal-Wallis test, followed (if Kruskal-Wallis significant) by pairwise Wilcox rank-sum tests with Bonferroni-Holm correction for multiple comparisons.

### Imaging & image analysis

We performed live-imaging on larvae embedded laterally in 1.2% low-melt agarose containing either 0.4% tricaine (still images) or 0.3 mg/ml pancuronium bromide (timelapse imaging) for immobilization. We acquired images using C-Apochromat 63x/1.20 NA water immersion objectives on a Zeiss LSM 880 and a Zeiss CellObserver Spinning Disk confocal system equipped with a Photometrics Prime 95B camera. We collected images of neutral red-stained larvae using a Zeiss AxioObserver microscope. After collecting images with Zen (Carl Zeiss), we performed all processing and analysis using Fiji/ImageJ. All image analysis was performed blind by using the Fiji plugin Lab-utility-plugins/blind-files.

### Microglia process motility analysis

Larvae were immobilized by immersion in pancuronium bromide 0.3 mg/ml, aided by a small incision at the tip of the tail, prior to mounting in agarose. Timelapse z-stack frames were acquired every 2.5 min for 1 hour. Movies were max-projected at every time frame, registered to correct for drift, and cells were manually tracked from frame to frame for process maintenance, withdrawal, or formation of new processes. Because spinal cord microglia at larval stages have very few processes (<10), events occurring at distalmost (usually primary or secondary) processes were tracked.

### Myelin inclusion measurements

Z-stack images of microglia, acquired with uniform optical sectioning and number of slices, were sum-projected in Fiji. The microglia border was manually traced, and the background was cleared. The channels were split and the myelin channel was subjected to morphological segmentation using the Fiji plugin MorphoLibJ (13). Specific parameters were: object image, gradient type=morphological, gradient radius=1px, watershed segmentation tolerance=60 (spinal cord) or 20 (optic tectum). Watershed segmentation values were predetermined on sample images of microglia from each of the two regions. Following segmentation, the wand tool was used to trace watershed ROIs and ROI and total cell area values were saved for downstream analysis in R.

### Sheath length measurements

Z-stack images of oligodendrocytes, acquired with uniform optical sectioning, were traced in 3D using the Fiji plugin Simple Neurite Tracer (19). The number of paths (sheaths) belonging to a single oligodendrocyte is the number of sheaths per cell and the path length in 3D is the sheath length. Default parameters were used, except cursor snapping = 2 XY pixels. Path lengths were exported to R for analysis.

## Supplemental Movies

**Movie S1. Engulfment of myelin sheaths by microglia.** Timelapse imaging (5 min/frame) of a microglia (yellow) interacting with an oligodendrocyte (magenta) in a *Tg(mpeg1.1:mVenus-CAAX; sox10:mRFP)* larva. Two myelin sheaths (magenta rectangles) are intact and visibly surrounded by microglial processes at first but become engulfed by the microglia soon after.

**Movie S2. 3D rotation of a microglia containing myelin inclusions.** A microglia (yellow) in a *Tg(mpeg1.1:mVenus-CAAX; mbpa:mCherry-CAAX)* larva containing phagocytosed myelin (magenta inclusions). Note intact myelin sheaths next to the microglia (magenta tubes). Movie generated with 3D Viewer plugin in Fiji.

## References and Notes

1. J. H. Hines, A. M. Ravanelli, R. Schwindt, E. K. Scott, B. Appel, Neuronal activity biases axon selection for myelination in vivo. Nat. Neurosci. 18, 683–689 (2015).

2. S. Mensch et al., Synaptic vesicle release regulates myelin sheath number of individual oligodendrocytes in vivo. Nat. Neurosci. 18, 628–30 (2015).

3. S. Koudelka et al., Individual Neuronal Subtypes Exhibit Diversity in CNS Myelination Mediated by Synaptic Vesicle Release. Curr. Biol. (2016), doi:10.1016/j.cub.2016.03.070.

4. M. Baraban, S. Koudelka, D. A. Lyons, Ca2+ activity signatures of myelin sheath formation and growth in vivo. Nat. Neurosci. 21, 19–23 (2018).

5. A. M. Krasnow, M. C. Ford, L. E. Valdivia, S. W. Wilson, D. Attwell, Regulation of developing myelin sheath elongation by oligodendrocyte calcium transients in vivo. Nat. Neurosci. 21, 24–28 (2018).

6. H. Wake et al., Nonsynaptic junctions on myelinating glia promote preferential myelination of electrically active axons. Nat. Commun. 6, 7844 (2015).

7. A. N. Hughes, B. Appel, Synaptic proteins expressed by oligodendrocytes mediate CNS myelination. bioRxiv, 410969 (2018).

8. P. Liu, J. Du, C. He, Developmental pruning of early-stage myelin segments during CNS myelination in vivo. Cell Res. 23, 962–964 (2013).

9. D. P. Schafer et al., Microglia Sculpt Postnatal Neural Circuits in an Activity and Complement-Dependent Manner. Neuron. 74, 691–705 (2012).

10. F. Ellett, L. Pase, J. W. Hayman, A. Andrianopoulos, G. J. Lieschke, mpeg1 promoter transgenes direct macrophage-lineage expression in zebrafish. Blood. 117 (2011) (available at http://www.bloodjournal.org/content/117/4/e49.full?sso-checked=true).

11. A. J. Svahn et al., Development of Ramified Microglia from Early Macrophages in the Zebrafish Optic Tectum. Inc. Dev. Neurobiol. 73, 60–71 (2012).

12. F. Peri, C. Nüsslein-Volhard, Live Imaging of Neuronal Degradation by Microglia Reveals a Role for v0-ATPase a1 in Phagosomal Fusion In Vivo. Cell. 133, 916–927 (2008).

13. D. Legland, I. Arganda-Carreras, P. Andrey, Bioinformatics, in press, doi:10.1093/bioinformatics/btw413.

14. J. K. Heppert et al., Comparative assessment of fluorescent proteins for in vivo imaging in an animal model system. Mol. Biol. Cell. 27, 3385–3394 (2016).

15. R. Almeida, D. Lyons, Oligodendrocyte Development in the Absence of Their Target Axons In Vivo. PLoS One. 11, e0164432 (2016).

16. J. R. Mathias, Z. Zhang, M. T. Saxena, J. S. Mumm, Enhanced cell-specific ablation in zebrafish using a triple mutant of Escherichia coli nitroreductase. Zebrafish. 11, 85–97 (2014).

17. P. Herbomel, B. Thisse, C. Thisse, Zebrafish Early Macrophages Colonize Cephalic Mesenchyme and Developing Brain, Retina, and Epidermis through a M-CSF Receptor-Dependent Invasive Process. Dev. Biol. 238, 274–288 (2001).

18. C. E. Shiau, K. R. Monk, W. Joo, W. S. Talbot, An Anti-inflammatory NOD-like Receptor Is Required for Microglia Development. Cell Rep. 5, 1342–1352 (2013).

19. M. H. Longair, D. A. Baker, J. D. Armstrong, Simple Neurite Tracer: open source software for reconstruction, visualization and analysis of neuronal processes. Bioinformatics. 27, 2453–2454 (2011).

20. N. A. Uranova, O. V Vikhreva, V. I. Rachmanova, D. D. Orlovskaya, Ultrastructural alterations of myelinated fibers and oligodendrocytes in the prefrontal cortex in schizophrenia: a postmortem morphometric study. Schizophr. Res. Treatment. 2011, 325789 (2011).

21. H. Neumann, M. R. Kotter, R. J. M. Franklin, Debris clearance by microglia: an essential link between degeneration and regeneration. Brain. 132, 288–95 (2009).

22. L. J. W. van der Laan et al., Macrophage phagocytosis of myelin in vitro determined by flow cytometry: phagocytosis is mediated by CR3 and induces production of tumor necrosis factor-a and nitric oxide. J. Neuroimmunol. 70, 145–152 (1996).

23. E. M. Gibson et al., Methotrexate Chemotherapy Induces Persistent Tri-glial Dysregulation that Underlies Chemotherapy-Related Cognitive Impairment. Cell. 176, 43–55.e13 (2019).

24. Q. Li et al., Developmental Heterogeneity of Microglia and Brain Myeloid Cells Revealed by Deep Single-Cell RNA Sequencing. Neuron. 101, 207–223.e10 (2019).

25. C. B. Kimmel, W. W. Ballard, S. R. Kimmel, B. Ullmann, T. F. Schilling, Stages of embryonic development of the zebrafish. Dev. Dyn. 203, 253–310 (1995).

26. K. M. Kwan et al., The Tol2kit: A multisite gateway-based construction Kit for Tol2 transposon transgenesis constructs. Dev. Dyn. 236, 3088–3099 (2007).

27. S. Curado, D. Y. R. Stainier, R. M. Anderson, Nitroreductase-mediated cell/tissue ablation in zebrafish: a spatially and temporally controlled ablation method with applications in developmental and regeneration studies. Nat. Protoc. 3, 948–54 (2008).

28. H. Wickham, Tidyverse: Easily install and load’tidyverse’packages. R Packag. version. 1 (2017).

29. C. O. Wilke, cowplot: streamlined plot theme and plot annotations for “ggplot2.” CRAN Repos (2016).

30. A. Kassambara, ggpubr:”ggplot2” based publication ready plots. R Packag. version 0.1. 6 (2017).

